# Functional Reorganization of the Somatomotor Network in Prodromal and Early Parkinson’s Disease

**DOI:** 10.1101/2025.09.29.679229

**Authors:** Adrian L. Asendorf, Verena Dzialas, Thilo van Eimeren, Merle C. Hoenig

## Abstract

In Parkinson’s disease (PD), higher network attack tolerance (NAT) may contribute to compensation of motor deficits. However, it is unclear whether NAT is lost due to disease progression or actively increased as a compensatory response.

We used cross-sectional resting state functional MRI data of 28 healthy controls (HC), 60 prodromal PD patients, 94 clinical PD patients to create graph theoretical networks. NAT was assessed at global and subnetwork level by calculating global efficiency upon iterative node removal. Using linear mixed-effects models we assessed how putaminal dopamine terminal (DaT) binding, or disease status affected NAT, controlling for density, age, sex and education. Finally, we compared the node degree distribution specifically for the somatomotor network (SMN) across groups.

Lower putaminal DaT predicted higher SMN NAT. Patients with PD showed elevated SMN NAT versus controls. Neither global nor other networks showed an effect. Compared to HCs subcortical/cerebellar SMN nodes appeared more connected in PD and prodromal patients.

Dopaminergic depletion appears to drive targeted reorganization of the SMN. This reorganization may involve additional recruitment of subcortical and cerebellar regions to sustain the information flow inside the SMN. Concomitantly, this active adaptation motivates further investigations regarding SMN NAT as potential compensation mechanism in early PD.

## Introduction

Over the past two decades, numerous studies have demonstrated that Parkinson’s disease is associated with widespread structural ^1^ and functional ^2^ alterations of brain connectivity. Notably, which of these alterations reflects passive consequences of neurodegeneration or active compensatory processes often remains vague. On a functional level, greater plasticity, adaptation, flexibility, and integration of certain brain networks could reflect compensation ^3–7^. However, quantifying such mechanisms remains challenging.

N*etwork attack tolerance* (NAT) is a graph-theoretical measure that has recently been associated with compensation. NAT captures how well a brain network can maintain effective information flow when crucial nodes of the network are iteratively removed ^8^. In PD, NAT has been associated with the preservation of both cognitive ^9^ and motor function ^10^, suggesting it may serve individual compensatory capacity. Indeed, we previously demonstrated that NAT of the somatomotor network (SMN) was positively associated with striatal dopamine transporter terminal (DaT) integrity, and motor function in early PD patients ^10^. However, while this study provides first indications that SMN NAT may actively serve as compensatory mechanism in clinical PD, it remains unknown whether it is also involved in the prodromal stage of PD, when significant dopaminergic loss is already present, but motor symptoms have not yet emerged. Hence, the dynamic of NAT in response to progressive dopaminergic loss across the disease course remains to be tested.

In this regard two opposing roles of NAT can be considered: 1) NAT as an inherent neurobiological trait that is progressively lost as PD advances. The extent of this loss can individually vary. 2) NAT as a mechanism that is actively increased in response to progressive dopaminergic decline – a process that likely begins during the preclinical stage of the disease.

To test these two assumptions, we utilized resting-state functional MRI (fMRI) and DaT imaging of a large, well-characterized cohort of 182 participants, including healthy controls (HC) and patients with prodromal (PM) and clinical PD. Across this sample, we investigated whether NAT of various large-scale networks, such as the SMN, shows a progressive decrease or increase with a reduction in DaT integrity (1) and across clinical disease stages (2). As our earlier study assessed only early PD (clinical symptoms <3 years), strong directional a priori predictions about NAT’s relation to disease stage or DaT integrity could not be justified. Therefore, analyses were exploratory.

## Materials and Methods

### Participants

This analysis included 182 participants, aged between 50 and 80 years, retrieved from the Parkinson’s Progression Markers Initiative (PPMI) in June 2025. Key inclusion criteria for the current study were: 1) available resting-state (rs) functional MRI (fMRI), structural MRI, and DaT single-photon emission computed tomography (DaT-SPECT), acquired within a time window of six months; 2) right-handedness; 3) no cognitive impairment (Montreal Cognitive Assessment ≥ 25); 4) no depressive symptoms (Geriatric Depression Scale ≤ 5), 5) no other neurological disorder; 6) availability of curated demographic data. All PD patients had an idiopathic PD diagnosis, a disease duration of less than three years, and had no dopaminergic therapy initiated (Levodopa equivalent daily dose (LEDD) = 0). The PM group comprised participants labeled with ‘prodromal synucleinopathy’ that had not received dopaminergic treatment yet (LEDD = 0). Healthy controls (HCs) were labeled ‘healthy control’. These selection criteria resulted in 28 HCs, 60 prodromal and 94 clinical PD patients. According to the Declaration of Helsinki all participants gave written consent, and each PPMI site obtained ethical approval for the study. Group demographics are summarized in Table 1. Additional details on selection procedures can be found in the supplements.

**Table 1:** Demographics of the study cohort. Standard deviation (SD) is provided for continuous variables and interquartile range (IQR) for ordinal values (MDS UPDRS-III, Education and MoCA scores). PD = Parkinson’s disease, PM= prodromal PD, HC = healthy control, DaT = dopamine transporter, MoCA = Montreal Cognitive Assessment, UPDRS-III = Movement Disorder Society Unified Parkinson’s Disease Rating Scale-Part III, *: p<.05

|  | HC (n=28) |  | PM (n=60) |  | PD (n=94) |  |  |
| --- | --- | --- | --- | --- | --- | --- | --- |
|  | Mean (SD) | MIN/MAX | Mean (SD) | MIN/MAX | Mean (SD) | MIN/MAX | p-value |
| Sex (m/f) | 16/12 | m=57% | 35/25 | m= 58% | 65/29 | m= 69% | .290 |
| Age | 66.7 (6.2) | 52.2 - 77.7 | 69.1 (5.6) | 52.9- 80.0 | 65.7 (7.1) | 50.4 - 79.8 | .013* |
| Education | 16 (2.5) | 7 - 20 | 17 (2) | 12 - 20 | 17(2) | 10 - 20 | .524 |
| MoCA | 28 (2) | 25- 30 | 28 (2) | 25 - 30 | 28 (3) | 25 - 30 | .686 |
| UPDRS III | 1.5 (2.2) | 0 - 7 | 3 (5) | 0 - 14 | 24 (16) | 8 - 57 | <.001* |
| Mean DaT (putamen) | 2.5 (0.4) | 1.88 - 3.46 | 1.7 (0.5) | 0.49 - 3.21 | 1.2 (0.4) | 0.16 - 2.07 | <.001* |

### fMRI acquisition and preprocessing

Since PPMI is a multi-center study, anatomical T1-weighted MRI and rs fMRI series were acquired on 32 different 3T Siemens scanners, comprising six models of the types: Prisma (34 subject IDs), Prisma fit (81 subject IDs), Verio (15 subject IDs), Skyra (41 subject IDs), Vida (10 subject IDs) and TrioTim (1 subject ID). The anatomical and rs-fMRI sequences followed the same specific protocol: Structural MRI: repetition time (TR) = 2300 ms, echo time (TE) = 1.9 -3 ms, flip angle = 9°, slice thickness = 1 mm, voxel size = 1x1x1 mm, number of slices = 256; Functional MRI: TR = 2500 ms, TE = 30 ms, flip angle = 80°, slice thickness = 3.5 mm, voxel size = 3.5x3.5x3.5 mm, number of volumes 240, acquisition time = approx. 10 min. Preprocessing of the fMRI data was carried out using CONN’s ^11^ default preprocessing pipeline (https://web.conn-toolbox.org/fmri-methods/preprocessing-pipeline), including spatial realignment, slice-timing correction, outlier identification, segmentation, normalization, and smoothing (Gaussian kernel, 7mm full-width half maximum).

### DaT-SPECT acquisition and preprocessing

As neurodegeneration in early PD is expected to mainly affect putaminal regions ^12,13^, only mean putamen values were retrieved from PPMI. DaT-SPECT ([^123^I]Ioflupane) imaging followed PPMI’s standard operating procedures (https://www.ppmi-info.org/study-design/research-documents-and-sops). Preprocessing involved raw projection reconstruction (HOSEM, HERMES), attenuation correction ^14^, 6 mm Gaussian filtering, and normalization to Montreal Neurological Institute space. Striatal binding ratios (SBRs) were calculated for the bilateral putamen using the occipital cortex as reference and average across left and right hemisphere.

### Graph theoretical network construction and network attack

Graph theory offers a powerful framework for modeling the brain as a network, where brain regions are represented as nodes and their functional connections as edges. Subsequently, a NAT analysis quantifies how efficiently a network continues to exchange information as its most connected and critical nodes are systematically removed.

In this analysis a set of 300 predefined functional regions of interest (ROIs) ^15^ were used to generate individual ROI-to-ROI connectivity matrices from the rs-fMRI data. These matrices served as the basis for constructing networks, which were then subjected to a targeted attack analysis. For each subject, this analysis estimated how well a network preserved its efficiency as nodes were removed. Nodes were removed iteratively in descending order of their degree of connectedness (i.e. the number of edges connected to this node). After each node removal the global efficiency of the network was calculated and plotted. Upon final node removal, the AUC of the resulting plot served as measure for NAT. Each subject received a NAT value, reflecting the percentage of maximum efficiency maintained during attacks. The procedure was repeated across nine network thresholds (densities), retaining 10–50% of the strongest connections (see Fig. 1A). This was done at the whole-brain (global) level as well as the single subnetwork level. The single subnetworks of interest comprised the default mode network (DMN, 65 ROIs), the frontoparietal network (FPN, 36 ROIs), the SMN (51 ROIs), and the attention network (ATN, 27 ROIs). For more detailed information on network construction and attack analysis, see Asendorf et al., (2025).

**Figure 1:**
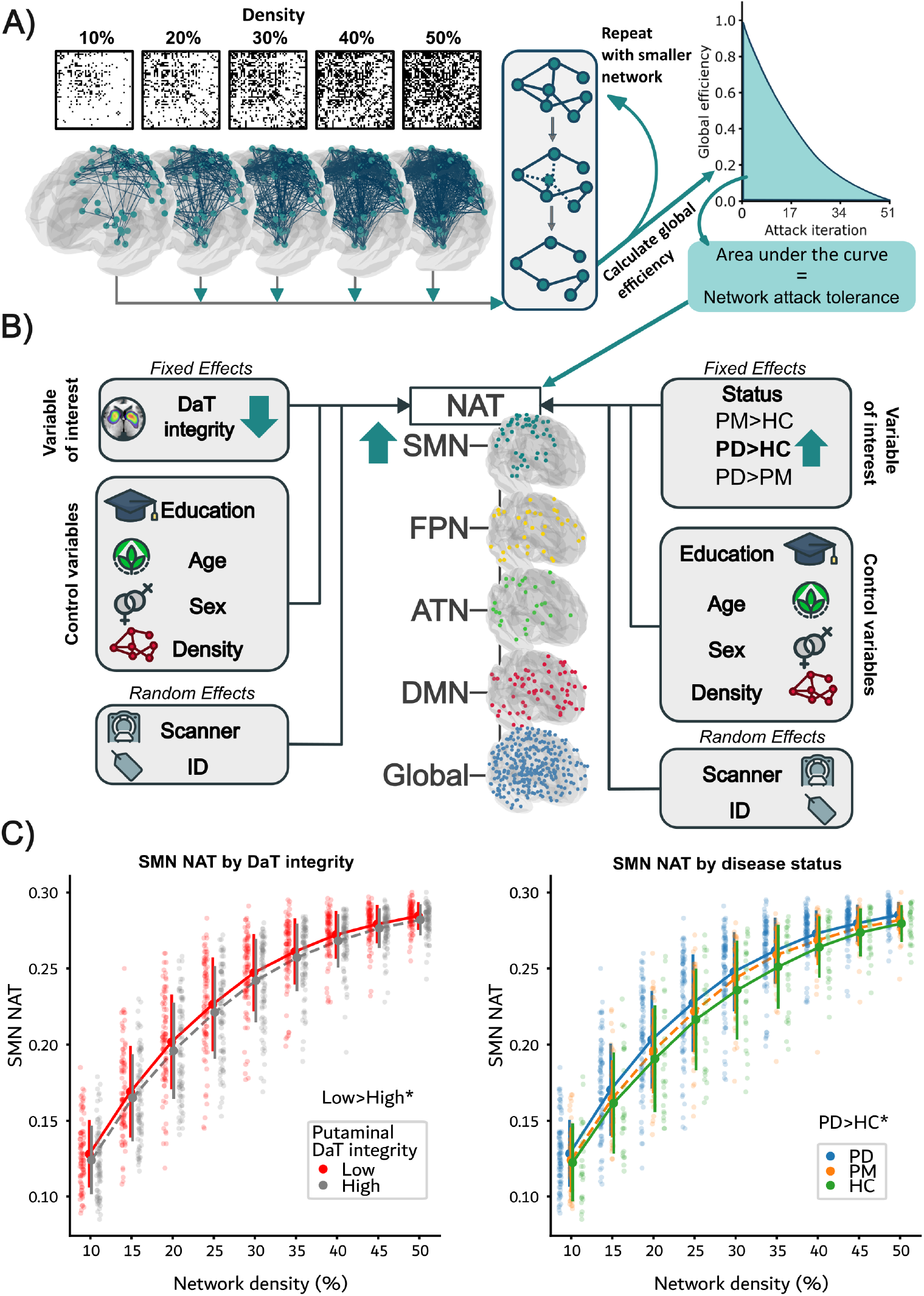
**A) Illustration of analysis pipeline**: **Left:** Region of interest (ROI) to ROI connectivity matrices were thresholded at nine different densities from 10% to 50% (for clarity only five out of nine densities are depicted), then binarized, and finally used for network construction. **Right:** During the attack analysis, the network’s nodes were iteratively removed in descending order of their connectedness. For each removed node, the network’s global efficiency was computed. The ability to sustain its global efficiency to successive attacks (area under the curve) defined the attack tolerance of the respective network. All analyses steps are illustrated using the SMN as an example. **B) Model structure and key results:** Left panel shows the effect of putamen dopamine terminal (DaT) integrity on NAT; right panel shows the effect of group status (Prodromal (PM), Parkinson’s disease (PD), and healthy controls (HC)) on NAT. The arrows are color-coded and represent the direction of the association between NAT of the respective network and the fixed effects. The surface projections depict the ROIs of the somatomotor network (SMN, turquoise), the frontoparietal network (FPN, yellow), the attention network (ATN, green), the default mode network (DMN, red), and the global network (blue). C**) SMN NAT plotted for each network density** and grouped by high and low (median split) putaminal DaT integrity (left) and disease status (right). Illustrations were plotted using nilearn ^29^, *: sign. Difference (p<.05) in linear mixed model

### Statistical analyses

To address our research goals, we first assessed the association between putaminal DaT integrity and global/subnetwork NAT across all individuals. Next, we compared NAT between the three groups. All models were controlled for network density, age, sex, and education. To account for the repeated measure design due to the different network densities, we performed two generalized linear mixed-effects models (GLMMs) ^16^, allowing crossed individual intercepts for each participant and MRI scanner. Fixed effects included: putaminal DaT-integrity or disease status, network density, age, sex and education. As disease status and putaminal DaT integrity were highly correlated with each other, we refrained from correcting our models for putaminal DaT integrity or disease status, respectively.

Since NAT is a fraction and by design never one or zero, a beta regression model was used ^17^, i.e., a GLMM with a beta response distribution utilizing a logit link function. Both statistical models were run separately for global and subnetwork NAT (5 runs = total of 10 models) using the ‘mgcv’ package in R (Rstudio ver. 2022.12.0). The significance level was set to p = 0.05. As our aim was to examine the presence or absence of associations between NAT and either dopamine availability or disease stage within each network independently, rather than to compare effects across networks, no correction for multiple comparisons was applied.

### Assessment of SMN node degree distribution

Given that the NAT analysis does not provide insights into the structural network alterations that underly shifts in NAT, we visualized the changes of SMN functional architecture across groups by using mean nodal degree maps. Mean nodal degree maps depict the average number of connections each of the 51 SMN ROIs had for each participant and were obtained by averaging the nodal degree for each ROI across the nine densities. To visualize general SMN network topology, these maps were averaged over all participants, mean centered and plotted. Next, to explore how functional architecture might change across groups, groupwise degree maps were averaged and used to plot pairwise group contrasts (e.g., PD–HC = mean PD – mean HC). Finally, to illustrate group differences in node degree distribution, frequency counts of node degrees for each group were smoothed into continuous probability distributions, using kernel density estimation.

## Results

### Effect of putaminal DaT integrity on NAT

GLMMs assessing the effect of putaminal DaT integrity on global/subnetwork NAT across all groups revealed a negative effect of mean putaminal DaT integrity on SMN NAT (β = -.030; p = .029). No significant associations were found for global NAT or the remaining subnetworks (Fig. 1B). For visualization purposes the raw SMN NAT was grouped by high and low (median split, 1,52 SBR) putaminal DaT integrity in Fig. 1C.

Regarding the covariates, network density displayed a robust positive relationship with NAT of every network. Further, higher age was linked to increased global, DMN and ATN NAT. Sex and education remained non-significant. Full model outputs are provided in Supplementary Table 2.

### Effect of disease status on NAT

The GLMMs testing the effect of disease status on global/subnetwork NAT, while controlling for demographic variables and network density, showed that patients with PD exhibited higher NAT within the SMN compared to HC (β = .059; p = .016; Fig. 1B). The PM group presented higher SMN NAT in comparison to the HC group, which did not reach significance (β = .034; p = .199). None of the other tested models resulted in significant group differences. Raw SMN NAT data plotted for each disease group can be found in Fig. 1C.

The covariates showed the same effects and directions as reported for the model assessing the effect putaminal DaT integrity on NAT. Importantly, in this model age was not linked to DMN NAT. All results are summarized in Supplementary Table 3.

### Qualitative comparison of SMN node degree distribution across groups

SMN nodes showed a clear topographical organization across the cohort: nodes with above-mean degree were predominantly located in cortical regions, whereas all subcortical and cerebellar nodes exhibited below-mean degree (Fig. 2A). Conversely, upon visual inspection, all subcortical and cerebellar nodes showed higher node degree values in the PD group in comparison to HCs. In PM patients, a high proportion of cerebellar and subcortical nodes had higher node degrees relative to HCs (Fig. 2B). In PD, nodes with extremely low and extremely high degrees occurred less, effectively increasing the probability of intermediate degree values (Fig. 2C).

**Figure 2:**
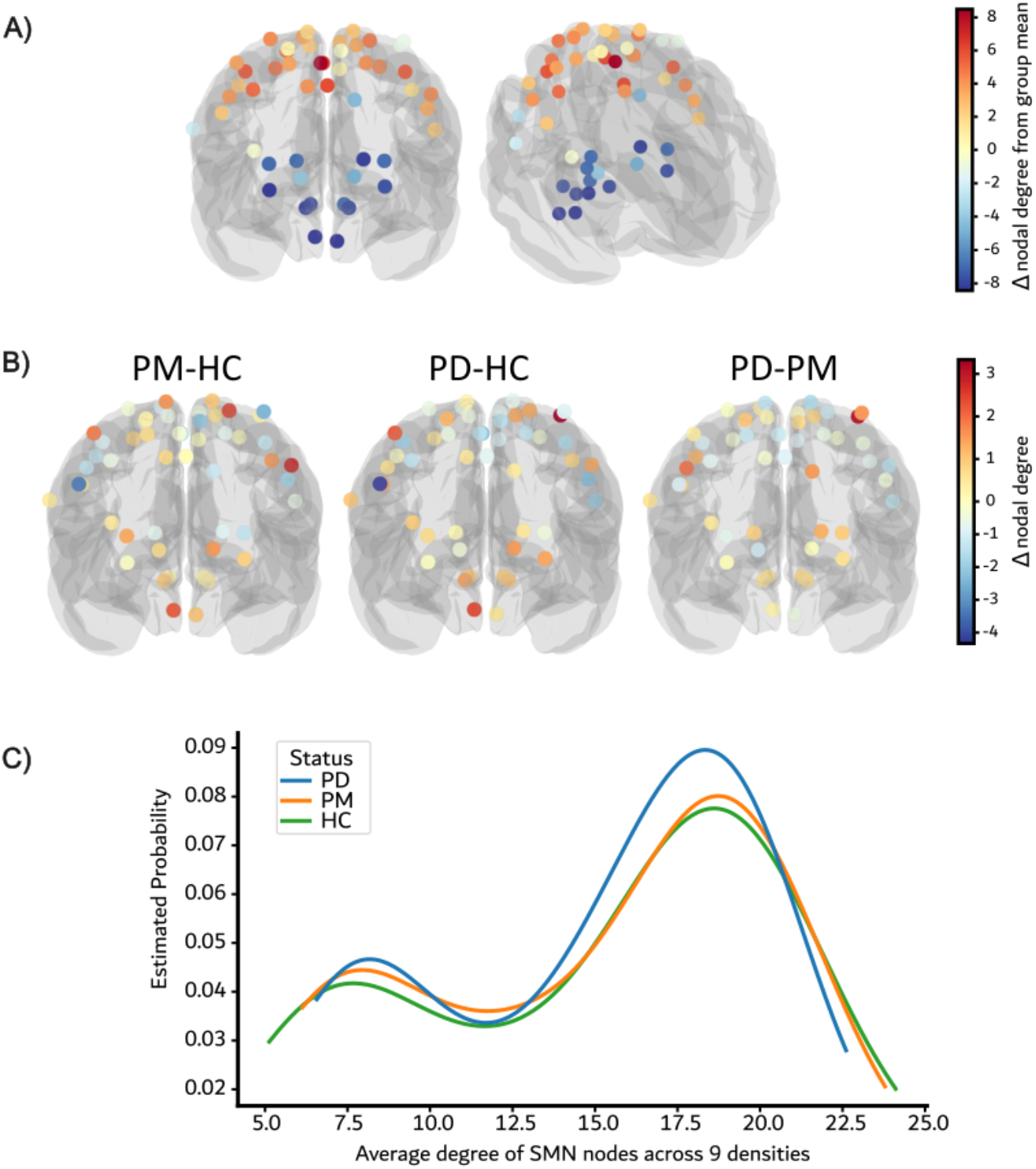
**(A) Average SMN node degree profile**. Surface renderings of the SMN (frontal, left; lateral, right). Node colors indicate mean-centered degree (node degree minus the SMN-wide mean), averaged across the entire cohort. Warm colors mark above-average degree; cool colors mark below-average degree. **(B) Group-level SMN node degree contrasts (PM, PD, HC)**. Frontal surface renderings of the SMN. Colors encode the difference for each node for the specified contrast (e.g., PD minus HC): warm = higher in the first group; cool = lower in the first group. **(C) Distributions of node degree across SMN by group (kernel density estimates)**. The y-axis reflects the estimated probability, with each group normalized independently to enable direct comparison of distribution shape. Each degree value represents the average across the nine density thresholds. Surface renderings were plotted using nilearn^29^

## Discussion

We investigated how dopaminergic terminal decline affects efficiency of brain networks after simulated node losses (i.e. NAT, network attack tolerance) before and after overt motor symptoms of PD. To this end, we used a large, well-characterized prodromal and clinical PD cohort from the PPMI database. Our primary finding was that SMN NAT increased with lower dopaminergic terminal availability. In other words, with increasing disease severity the SMN became more resilient to simulated attacks, suggesting that the functional architecture of this network undergoes a reorganization when the system becomes impacted by higher pathological load. This finding was further supported by the observation that PD patients exhibited increased NAT in the SMN compared to HCs. In PM individuals, SMN NAT did not differ from that of either HCs or clinical PD patients, but PM data occupied the space between PD and HC.

Importantly, even though we tested NAT in several subnetworks, only the SMN showed a disease-related effect, while other networks (i.e. DMN, FPN, ATN and whole brain) remained unaffected. This is consistent with the current understanding of PD pathophysiology: The early nigrostriatal degeneration primarily disrupts basal ganglia and, subsequently, cortical motor circuits ^18^. Prior PD imaging studies reported that especially the SMN and the basal ganglia together with the DMN appeared to be affected by functional changes associated with disease progression ^19^. Importantly, various studies specifically suggested functional changes related to dopamine medication intake ^20,21^ or striatal dopamine availability ^22–24^ in regions involving key nodes of our SMN network. Our findings align with these reports indicating that the architecture of the SMN changes as dopamine terminal availability declines.

Notably, while several studies have already reported a functional reorganization of the SMN in PD, it nonetheless remains unknown which exact functional changes represent compensational rather than maladaptive pathological mechanisms ^25^. Recent studies linked specific functional connectivity alterations to motor compensation ^23,24^. While these findings form a patchwork of hyper- and hypo-connectivity states and are thus hard to draw conclusions from, the assessment of NAT approximates real-world pathology propagation by sequentially removing network nodes, thereby revealing the networks’ ability to cope with loss of critical hubs. Prior evidence from an early PD cohort, suggests that increased SMN NAT reflects motor reserve ^10^. Extending this, our current findings indicate that dopamine depletion may drive a targeted reorganization of the SMN. Visual inspection of the degree distribution within the SMN revealed increased connectivity in typically lower-degree subcortical and cerebellar nodes in PM and PD patients. This is consistent with recent reports of additional subcortical and motor region recruitment in motor compensation ^26,27^. These findings overall suggest that when striatocortical input is disrupted, the SMN reroutes information through alternative, lesser involved pathways to preserve its function. In parallel, the proportion of nodes with intermediate degrees appeared increased in the current study, while that of nodes with extremely high or low degrees decreased in the PD cohort. This redistribution may reduce vulnerability to the loss of highly connected hubs, allowing the SMN to preserve efficiency despite the loss of critical cortical hubs.

Notably, this change in network structure may already begin in prodromal stages of the disease. In the current analysis no significant differences in terms of SMN NAT were observed between PM and HC. Yet, a minor positive effect was observable across all networks densities in addition to changes in degree distribution within the SMN in the PM group. The lack of significance may be due to subtle and below-threshold changes that cannot yet be statistically detected.

Importantly, the current assessments were limited by scanner type variability and their cross-sectional nature. Since DaT-SPECT has only recently been proposed as marker of PD progression ^28^, validation in longitudinal designs is warranted. Additionally, more refined motor tests, which are sensible to subtle motor changes in PM patients, could determine whether NAT plays a role in the phenoconversion from prodromal to manifest PD.

In conclusion, our findings indicate that striatal DaT loss is associated with a targeted reorganization of the SMN, enabling it to sustain efficient information flow despite the loss of critical cortical hubs. This suggests that, in clinical PD, the brain actively adapts to progressive striatal dopaminergic degeneration. However, its relevance to motor function needs to be further validated in longitudinal approaches.

## Data availability statement

Data used in the preparation of this article was obtained in June 2025 from the Parkinson’s Progression Markers Initiative (PPMI) database (www.ppmi-info.org/access-dataspecimens/download-data), RRID:SCR_006431. For up-to-date information on the study, visit www.ppmi-info.org.

