## SupplementaryMaterial for "Functional Reorganization of the Somatomotor Network in Prodromal and Early Parkinson’s Disease"

### Supplements:

#### PPMI data extraction and curation

Data were obtained from the Parkinson's Progression Markers Initiative (PPMI) in June 2025 via the Image and Data Archive (IDA), selecting participants aged between 50–80 years from the three respective groups: PD, PM, and HC groups. We included the following visits: Screening, Baseline, and Months 3, 6, 9, 12, 18, and 24. Filters were applied for radiopharmaceutical use (DAT) and MRI/fMRI data acquired on Siemens scanners. This yielded metadata for 2078 subjects. The resulting CSV file was cleaned to correct modality labels, reformat date-time fields, and replace SPECT entries with higher-resolution DaT-SPECT metadata from PPMI spreadsheets (September 2024; PM subset July 2023). Only SPECT sessions within ±6 months of MRI scans were retained. Demographic data was added using the "curated data set" (March 2025). Next, filtering criteria outlined in table 2 were applied globally and at the group level (see Table 2), resulting in 399 subjects: 131 PM, 223 PD, and 45 HC. Respective imaging data was downloaded via IDA and converted from DICOM to NIfTI using dcm2niix (v1.0.20211006). Using the ".json" files we assured that all MRI and fMRI sessions followed one identical sequence, reducing the sample to 213 subjects. Visual inspection excluded 10 cases (ghosting, structural lesions). Following preprocessing, automated quality control excluded 21 more (signal drift, motion artifacts), resulting in a final analytic sample of 182 participants.

| | |
|---|---|
| Whole group: | ✓ MRI + fMRI + DaT SPECT available<br>✓ Time delta between fMRI and DaT SPECT ≤ 6 months<br>✓ Curated data available<br>✓ Absence of cognitive impairment: MoCA ≥ 25<br>✓ Age between: 50 -80<br>✓ Absence of depressive symptoms: GDS ≤ 5<br>✓ Only right-handed participants<br>✓ Absence of any other neurological condition |
| PD group | ✓ Disease duration < 3 Years<br>✓ Unmedicated (LEDD = 0mg) |

| | |
|---|---|
| | ✓ Primary diagnosis of 'idiopathic PD' |
| PM group | ✓ Only Baseline scans<br>✓ Unmedicated (LEDD = 0mg)<br>✓ Primary diagnosis of 'Prodromal Synucleinopathy' |
| Control group | ✓ Primary diagnosis of 'Healthy Control' |

**Supp. Table 1. Inclusion criteria applied to the IDA advanced search result:** Whole-group criteria were applied first, ensuring that every participant (PD, PM, HC) had complete multimodal imaging (MRI, resting-state fMRI, and DaT SPECT) acquired within six months of each other, met basic demographic and health requirements (age 50–80 y, right-handed, MoCA ≥26, GDS ≤5, no other neurological disorders), and had fully curated clinical records at PPMI. Group-specific criteria were then layered on: idiopathic Parkinson's disease (PD) within three years of diagnosis and unmedicated (LEDD = 0 mg); prodromal synucleinopathy (PM) at baseline, also unmedicated; and healthy controls (HC) without neurological or psychiatric diagnoses. DaT = dopamine transporter, GDS = Geriatric Depression Scale, MoCA = Montreal Cognitive Assessment, UPDRS-III = Movement Disorder Society Unified Parkinson's Disease Rating Scale-Part III, LEDD = Levodopa equivalent daily dose

| Fixed effects | Global | | | | SMN | | | | DMN | | | | ATN | | | | FPN | | | |
|---|---|---|---|---|---|---|---|---|---|---|---|---|---|---|---|---|---|---|---|---|
| | ß | *p* | CI | | ß | *p* | CI | | ß | *p* | CI | | ß | *p* | CI | | ß | *p* | CI | |
| Biological factor | | | | | | | | | | | | | | | | | | | | |
| Mean putamen | -.001 | .807 | .-.005 | .004 | **-.030** | **.029*** | **-.057** | **-.003** | .013 | .276 | .-.010 | .035 | .008 | .457 | -.013 | .028 | -.009 | .375 | -.028 | .011 |
| Control variables | | | | | | | | | | | | | | | | | | | | |
| Age | **.001** | **.009*** | **.000** | **.001** | -.001 | .679 | -.002 | .003 | **.002** | **.034*** | **.000** | **.004** | **.002** | **.025*** | **.000** | **.004** | .001 | .532 | -.001 | .002 |
| Sex(f<m) | .003 | .278 | -.002 | .009 | -.015 | .401 | .-.050 | .020 | .025 | .106 | -.005 | .055 | .017 | .226 | -.010 | .043 | .005 | .725 | -.021 | .030 |
| Education | .000 | .729 | -.001 | .001 | .002 | .566 | -.009 | .005 | -.001 | .712 | -.007 | .005 | .001 | .599 | -.004 | .007 | .000 | .800 | -.006 | .004 |
| Network density | **.106** | **<.001*** | **.104** | **.109** | **.616** | **<.001*** | **.606** | **.625** | **.542** | **<.001*** | **.0533** | **.550** | **.747** | **<.001*** | **.735** | **.760** | **.676** | **<.001*** | **.666** | **.685** |
| Explained var. | $R^2$=.81 N=198 | | | | $R^2$=.93 N=198 | | | | $R^2$=.91 N=198 | | | | $R^2$=.91 N=198 | | | | $R^2$=.94 N=198 | | | |

**Supp. Table 2: Results table of GLMMs assessing contribution of DaT integrity in the putamen to network attack tolerance (NAT) in five different networks.** Significant findings are highlighted in bold. ß = unstandardized beta coefficients, *: p<.05, CI represents the 95% confidence interval, ATN = attention network, DMN = default mode network, FPN = frontoparietal network, SMN =somatomotor network.

| Fixed effects | Global | | | | SMN | | | | DMN | | | | ATN | | | | FPN | | | |
|---|---|---|---|---|---|---|---|---|---|---|---|---|---|---|---|---|---|---|---|---|
| | ß | *p* | CI | | ß | *p* | CI | | ß | *p* | CI | | ß | *p* | CI | | ß | *p* | CI | |
| Variable of Interest | | | | | | | | | | | | | | | | | | | | |
| Status (PD > HC) | .002 | .619 | *-.006* | *.010* | **.059** | **.016*** | ***-.018*** | ***.087*** | .015 | .488 | *-.027* | *.056* | -.008 | .667 | *-.045* | *.029* | .014 | .344 | *-.021* | *.049* |
| Status (PM > HC) | .001 | .791 | *-.007* | *.009* | .034 | .199 | *.011* | *.107* | .035 | .135 | *-.011* | *.080* | -.018 | .361 | *-.057* | *.021* | .018 | .425 | *-.019* | *.056* |
| Status (PD > PM) | .001 | .802 | *-.005* | *.007* | .025 | .218 | *-.015* | *.064* | -.020 | .250 | *.-.054* | *.014* | .010 | .489 | *-.019* | *.039* | -.004 | .789 | *-.031* | *.024* |
| Control variables | | | | | | | | | | | | | | | | | | | | |
| Age | **.001** | **.009*** | ***.000*** | ***.001*** | .001 | .400 | *-.001* | *.004* | .002 | .087 | *.000* | *.004* | **.002** | **.020*** | ***.000*** | ***.004*** | .001 | .558 | *-.001* | *.002* |
| Sex(f<m) | .003 | .267 | *-.002* | *.009* | -.011 | .540 | *-.045* | *.024* | .021 | .154 | *-.008* | *.051* | .014 | .302 | *-.012* | *.040* | .007 | .598 | *-.018* | *.032* |
| Education | .000 | .732 | *-.001* | *.001* | -.002 | .637 | *-.009* | *.005* | -.002 | .548 | *.-.008* | *.004* | .001 | .623 | *-.004* | *.007* | -.001 | .830 | *-.006* | *.004* |
| Density | **.107** | **<.001*** | ***.104*** | ***.109*** | **.616** | **<.001*** | ***.606*** | ***.625*** | **.542** | **<.001*** | ***.533*** | ***.550*** | **.747** | **<.001*** | ***.735*** | ***.760*** | **.676** | **<.001*** | ***.666*** | ***.685*** |
| Explained var. | $R^2$=.81. N=1638 | | | | $R^2$=.92. N=1638 | | | | $R^2$=.92. N=1638 | | | | $R^2$=.91 N=1638 | | | | $R^2$=.94 N=1638 | | | |

**Supp. Table 3: Results table of GLMMs assessing contribution of disease status to network attack tolerance (NAT) in five different networks.** Significant findings are highlighted in bold. ß = unstandardized beta coefficients, *: p<.05, CI represents the 95% confidence interval, PD= Parkinsons disease, HC = healthy control, PM prodromal, ATN = attention network, DMN = default mode network, FPN = frontoparietal network, SMN =somatomotor networ
